# Virility does not Imply Immensity: Testis size, Accessory Gland Size and Ejaculate depletion pattern do not Evolve in Response to Experimental Manipulation of Sex Ratio in *Drosophila melanogaster*

**DOI:** 10.1101/032649

**Authors:** Tejinder Singh Chechi, Syed Zeeshan Ali, Nagaraj Guru Prasad

## Introduction

Females in many species mate more than once and store sperm from more than one male. This leads to post-copulatory competition where sperm from different males compete to fertilize the limited number of eggs produced by the female-typically called Sperm competition (Parker, 1970a) (Wedell et al., 2002). Sperm competition and the resulting post-copulatory selection can significantly alter male reproductive behavior (Simmons et al., 1993) (Cook & Wedell, 1996) (Gage and Barnard, 1996) (Wedell & Cook 1999a,b) (Bretman et al., 2009) (Bretman et al., 2010) (Nandy & Prasad, 2011) and physiology (Wolfner 1997). Since a large number of sexual species are promiscuous, sperm competition is expected to be a widespread phenomenon influencing the evolution of male anatomy, physiology and behaviour.

There are at least two ways in which males can increase the probability of their success in sperm competition. One way is to increase the production of ejaculate. Sperm competition theory (Parker, 1970a) predicts that when competition is high, selection favors increased reproductive investment. A male that produces and transfers more number of sperm is likely to sire more progeny under competitive conditions. Positive correlation between sperm number and sperm competitive ability has been reported (Snook 2005). Further, increase in testes size often results in increased sperm number. (Stockley et al., 1997) found increase in gonad size and sperm number with increase in intensity of sperm competition across species of fishes. Promiscuous mammalian species have larger testes which produce more sperms (Kenagy & Trombulak, 1986). Similarly, a direct correlation between sperm competition and gonad size has been established in various organisms like fishes (Stockley et al., 1997), butterflies (Gage, 1994) and yellow dung flies (Hosken & Ward, 2001).

In some species like *D. melanogaster*, in addition to sperms, other components of the ejaculate like the Accessory gland proteins (Acps) are known to play an important role in sperm competition. Along with facilitating sperm transfer, Acps exert wide-ranging effects on female reproductive behavior and improve male's chances of siring a significant proportion of the female’s offspring. Acps affect many aspects of the female’s reproductive activity and behaviour (Aigaki et al., 1991) (Wolfner, 2009). The Acps may render her unwilling to remate for some time (Chapman, 2001), induce her to ovulate by stimulating octopaminergic signaling (Frank W. Avila et al., 2012) etc. These effects on the behavior of the mated female are often long lasting (McGlaughon & Wolfner, 2013), thus ensuring that any eggs laid will be fertilized by that male’s sperm. Thus, it is quite possible that in such species, increased sperm competition can select for increased Acp production along with an increase in sperm production, leading to an increase in accessory gland size.

Another way for males to increase their chances in sperm competition is to strategically invest the ejaculate. Ejaculate production is costly for males (Dewsbury, 1982) and ejaculate transferred during mating is an important determinant of the outcome of sperm competition (Moller, 1987). Therefore, one would expect males to plastically vary the amount of ejaculate transferred based on their perception of sperm competition risk. (Bretman et al., 2009) show that males alter their ejaculate investment (measured as mating duration) according to the level of sperm competition. (Nandy & Prasad 2011) show that *Drosophila melanogaster* males can plastically vary copulation duration based on their perception of sperm competition. Such variation in copulation duration is also positively correlated with sperm defense ability.

(Linklater et al. 2007) studied gonad size and ejaculate depletion pattern in *D. melanogaster* males from populations evolving under male biased and female biased operational sex ratios. Compared to flies maintained under female biased sex ratio, flies maintained under male biased operational sex ratio experience higher male-male competition and show increased number of mating per female. Thus the intensity and risk of sperm competition is expected to be higher in male biased populations. Therefore, altered operational sex ratios can potentially lead to the evolution of sperm competitive ability, ejaculate size and/or gonad size. (Linklater et al., 2007) found that virgin males from the two regimes showed no difference in either testes size or accessory gland size. However, when allowed to mate, males from the male-biased regime depleted their Acps at a much faster rate than the males from the female-biased regime, as measured by the reduction in accessory gland size upon mating.

We maintained replicate populations of *D. melanogaster* under male-biased (M), female-biased (F) and equal (C) sex ratios for over 100 generations. In an earlier study (Nandy et al., 2013) found that males from M populations had evolved increased sperm defense and offense ability compared to the males from F populations. In the present study, address the mechanistic basis of the increased sperm competitive ability of M population males. We specifically ask the following questions-

a. Does the size of the testes and/or accessory glands evolve in response to altered levels of sperm competition?
b. Does the ejaculate investment pattern evolve in response to altered levels of sperm competition? Following (linklater et al., 2007) we use the change in the testes/accessory gland area post mating as a measure of ejaculate investment.

## Materials and Methods

### Maintenance of population

In the present study we used the ‘M’ and ‘F’ populations of *D. melanogaster* as described in (Nandy et al., 2013). The selected populations were derived from a long-term laboratory adapted population of *D. melanogaster*, LHst (Prasad et al., 2007). LHst is a derivative of LH base population (see Chippindale et al., 2001 for details of the maintenance regime) having a recessive autosomal marker “scarlet eye”. The male biased sex ratio regime (M) had a ratio of three males to one female while the female biased reime (F) had a ratio of one male to three females. Each regime had three replicate populations (M1–3 and F1–3) (see Nandy et al., 2013 for details of selection history). The populations are maintained on a 14-day discrete generation cycle, at 25°C and 60% relative humidity (RH) and 12:12 hours light/dark cycle. They are fed on standard cornmeal– molasses–yeast food in standard vials (90-mm length × 30-mm diameter). Flies are grown in controlled larval density of 140– 160 per vial. We collect virgin flies each generation and hold them in single sex vials (eight individuals per vial) till the 12th day after egg collection. They are then combined following the respective sex ratio regimes in food vials provisioned with 0.47 mg per female of live yeast (adult competition vials). Two days later, flies are transferred to new vials for oviposition. The eggs laid during the window of 18hrs of oviposition are used to start the next generation.

To account for any nongenetic parental effects, all the populations are passed through one generation of common rearing conditions during which all the populations are maintained at 1:1 sex ratio. We call this process as “standardization” (For details see Nandy et al., 2013). The flies for the assays were generated from standardized populations.

### Generation of experimental flies

Males from M (male-baised) and F (female-baised) selection regimes were used for the experiment. For the mating trials, common females from Lhst (base-line population) were used. We generated all the experimental flies under controlled larval density and standard culture conditions (25°C, 60–80% RH, 12 hours–12 hours light / dark cycle). 150 eggs were cultured in 8–10 mL of cornmeal-molasses food per vial, for each of the seven selected populations (2 selection regimes X 3 replicates, 1 LHst). On 10th day after egg collection, adult flies started emerging. We collected M and F males as virgins by isolating them within 6 hours of eclosion, under light CO_2_ anesthesia. These males were held in single sex vials, at the density of 8 per vial for 2 days. LHst females were collected and maintained in the similar manner as described above.

### Experimental design

We had two sets of mating treatments for the experiment On day 12 post egg collection (2 to 3 days post eclosion), males collected from M and F populations were randomly assigned to one of the two mating treatments (a) virgin males and (b) mated only once. Males from the virgin treatment (10 single-sex vials with virgin M or F males) were flash frozen. For the mated treatment, 8 virgin males from M or F populations were combined with 10 LHst virgin females (to ensure all males mate once and only once). 10 such vials were set up per selection regime × replicate population combination. The flies were allowed to mate once. Single mating was ensured by manual observation. Typically, mating pairs are formed within 5 minutes of combining the males and females. Mating typically lasts for about 20 minutes. When the last mating pair decoupled in a vial, all the flies in the vial were flash-frozen. Thus we had two treatments for each selection regime: Virgin and Singly mated.

## Dissections and Measurements

The frozen flies were dissected in 9μL 1X-PBS solution. The testes, accessory glands and thorax were imaged using the ZEISS AxioCam ICc 1, attached to Zeiss Stemi 2000-C stereozoom microscope.

All thorax images were taken at 3.5 X magnification. Thorax length was calculated between two fixed points (Figure 1) using the length tool in Image J software (National Institutes of Health, http://imagej.nih.gov/ij/). Each thorax was measured thrice and the average of these three values was used as a measure of thorax length. Testes and accessory glands were imaged at 5 X magnification (Figure 2 and Figure 3). Testes area and accessory gland area were measured using Image J. The area measured was the cross section area as seen in the image. The measurements required combination of different tools. The brightness and contrast values for each image were adjusted to bring out the subject of the picture (Testes/accessory glands) from the background. Using the Threshold adjustment, the area of the subject was detected by the software. Then using the wand tool, the area to be measured was marked and measured (Figure 4 and Figure 5). All measurements were repeated thrice and an average of these three measurements was used as a measure of the area. We measured the right and left testes and accessory glands as mentioned above. We then averaged the areas of the right and left testes (or accessory glands) to get an average testes (or accessory gland) size.

**Figure 1:**
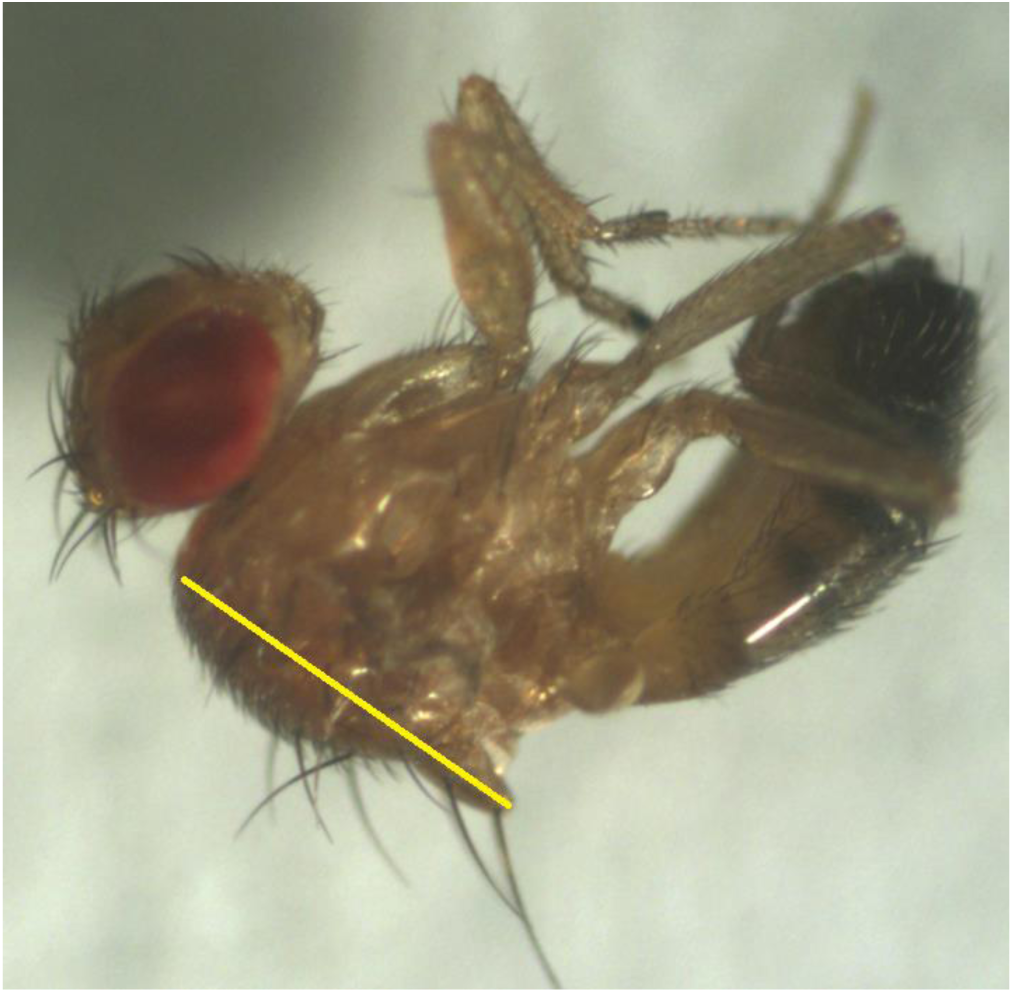
Thorax length measured between two fixed points at 3.5X magnification and 1X objective.

**Figure 2:**
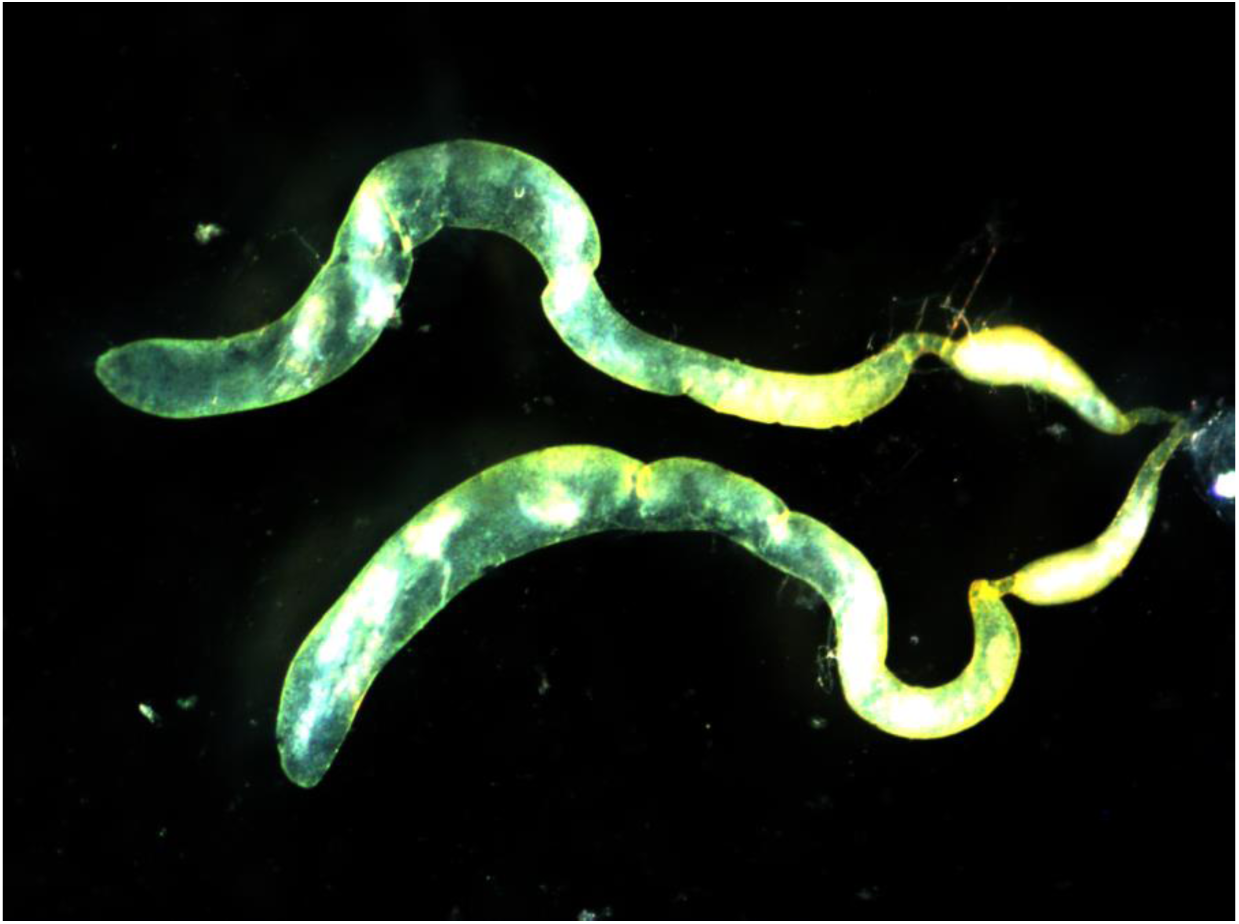
Image of pair of testes dissected out from a male *Drosophila melanogaster*, for measurement.

**Figure 3:**
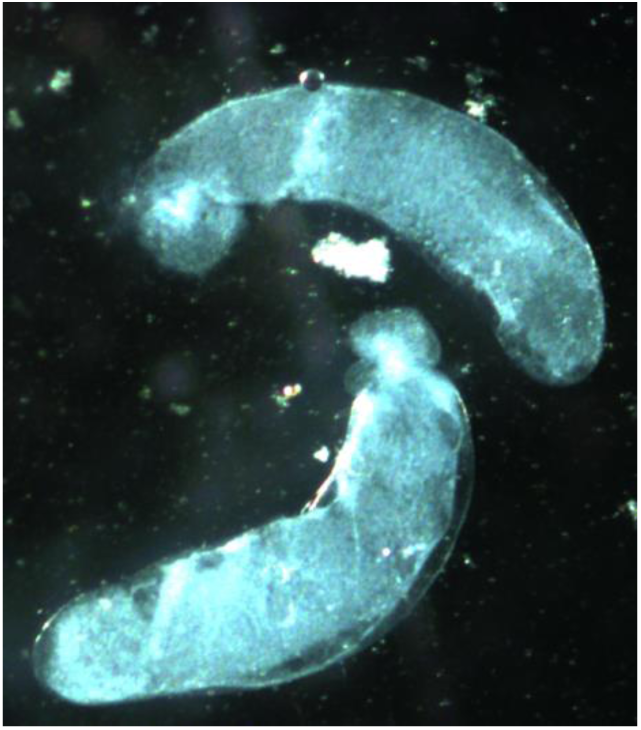
Image of pair of accessory glands dissected out from male *Drosophila melanogaster*, for measurement.

**Figure 4:**
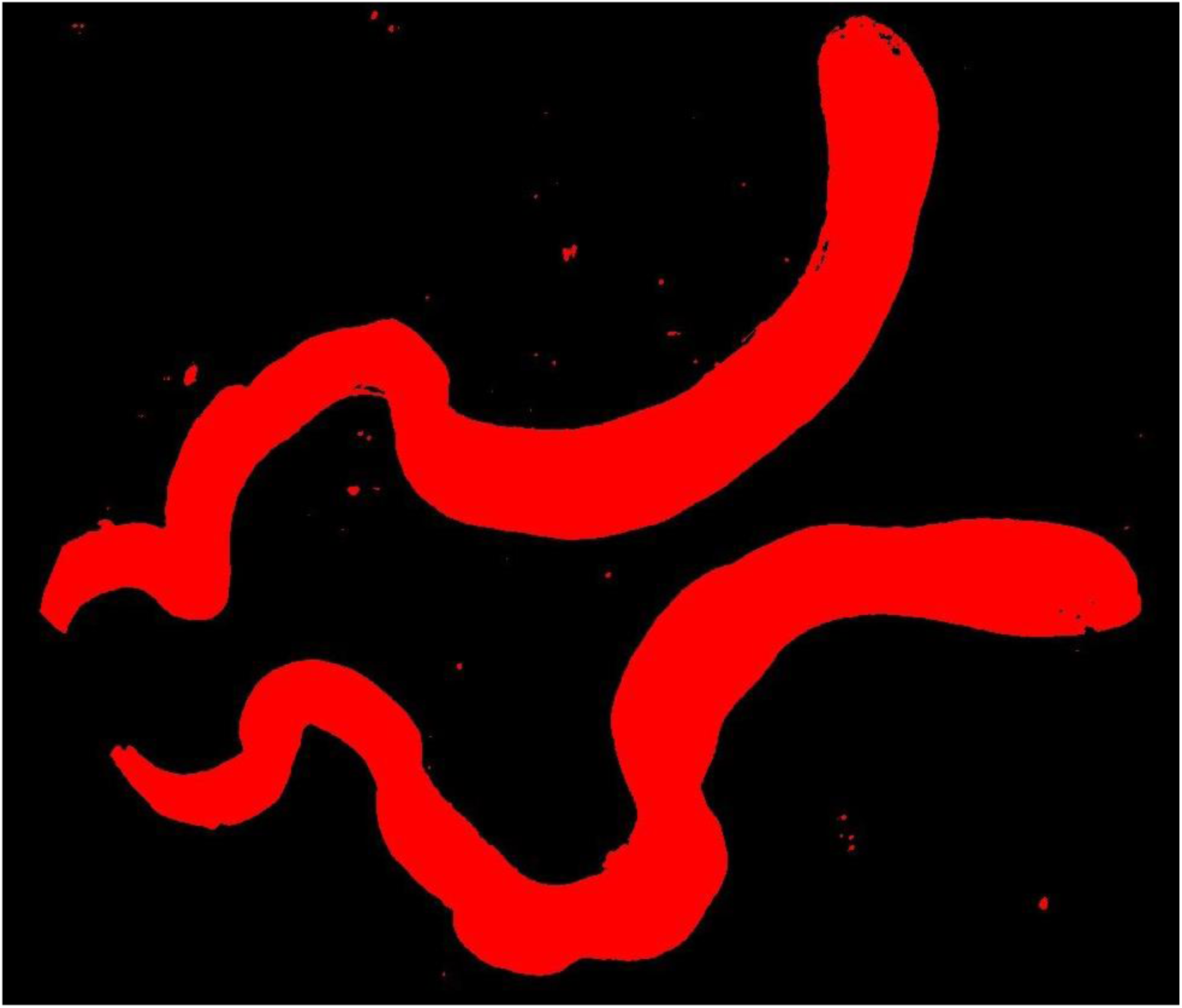
Selected area of pair of testes for measurement.

**Figure 5:**
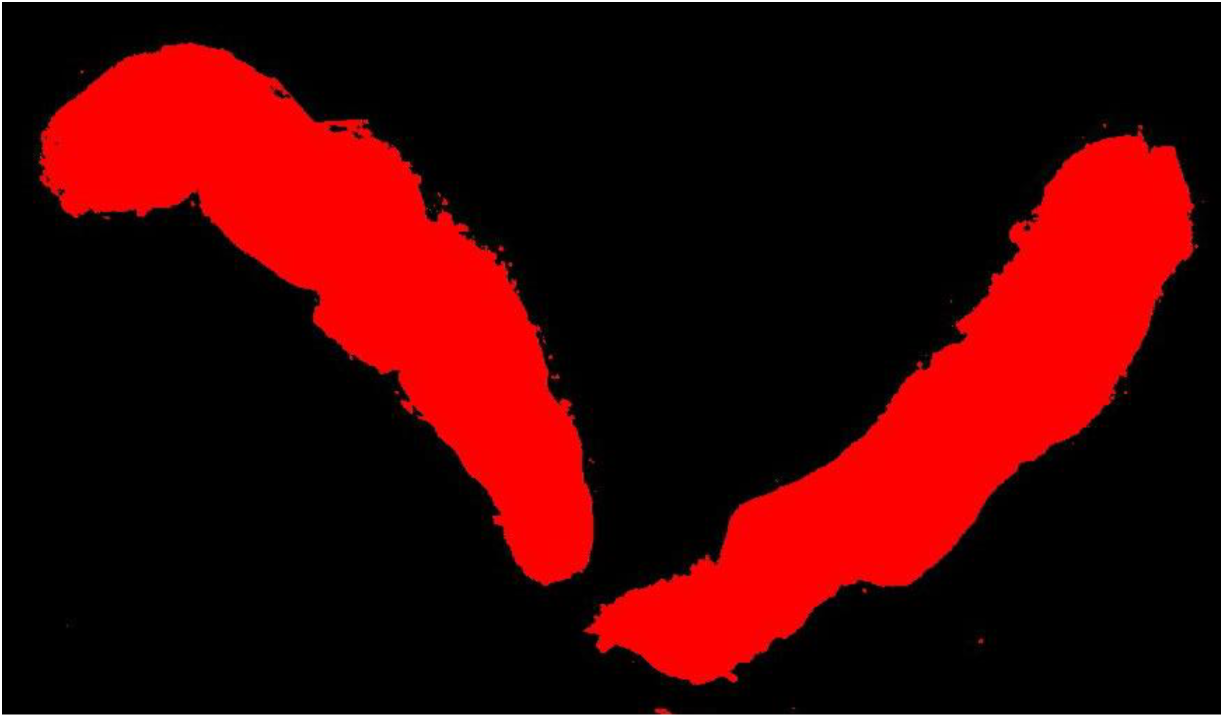
Selected area of pair of accessory glands for measurement.

## Results

We calculated the mean testes area and accessory gland area for each fly from the pair of testes and accessory glands respectively. The testes area and accessory gland area was standardized to the body size by dividing it with the thorax length measure of the same fly. Then the standardized testes area and standardized accessory gland area were analyzed using mixed model analysis of variance (ANOVA), with selection regime and mating status as fixed factors crossed amongst themselves and with random blocks.

We found no significant effect of selection regime or mating status on testes area (Table 1). The testes area for the virgin males as well as that of the mated males were not significantly different from each other. (Figure 6). Taken together, these results indicate that neither selection history nor immediate mating status affect testes size.

**Table 1:** Multivariable mixed model analysis of variance (ANOVA) on testes cross section area with block as random factor and selection regime and mating treatments as fixed factors.

| Source | df num | Ddf den | f-ratio | p-value |
| --- | --- | --- | --- | --- |
| Sel | 1 | 2 | 0.02 | 0.907 |
| Mating Status | 1 | 2 | 0.06 | 0.829 |
| Block | 2 | 3.5 | 0.59 | 0.6 |
| Sel*Mating Status | 1 | 2 | 2.32 | 0.267 |
| Sel*Block | 2 | 2 | 25.68 | 0.037 |
| Mating Status*Block | 2 | 2 | 52.53 | 0.019 |
| Sel*Mating Status*Block | 2 | 287 | 0.38 | 0.683 |

**Figure 6:**
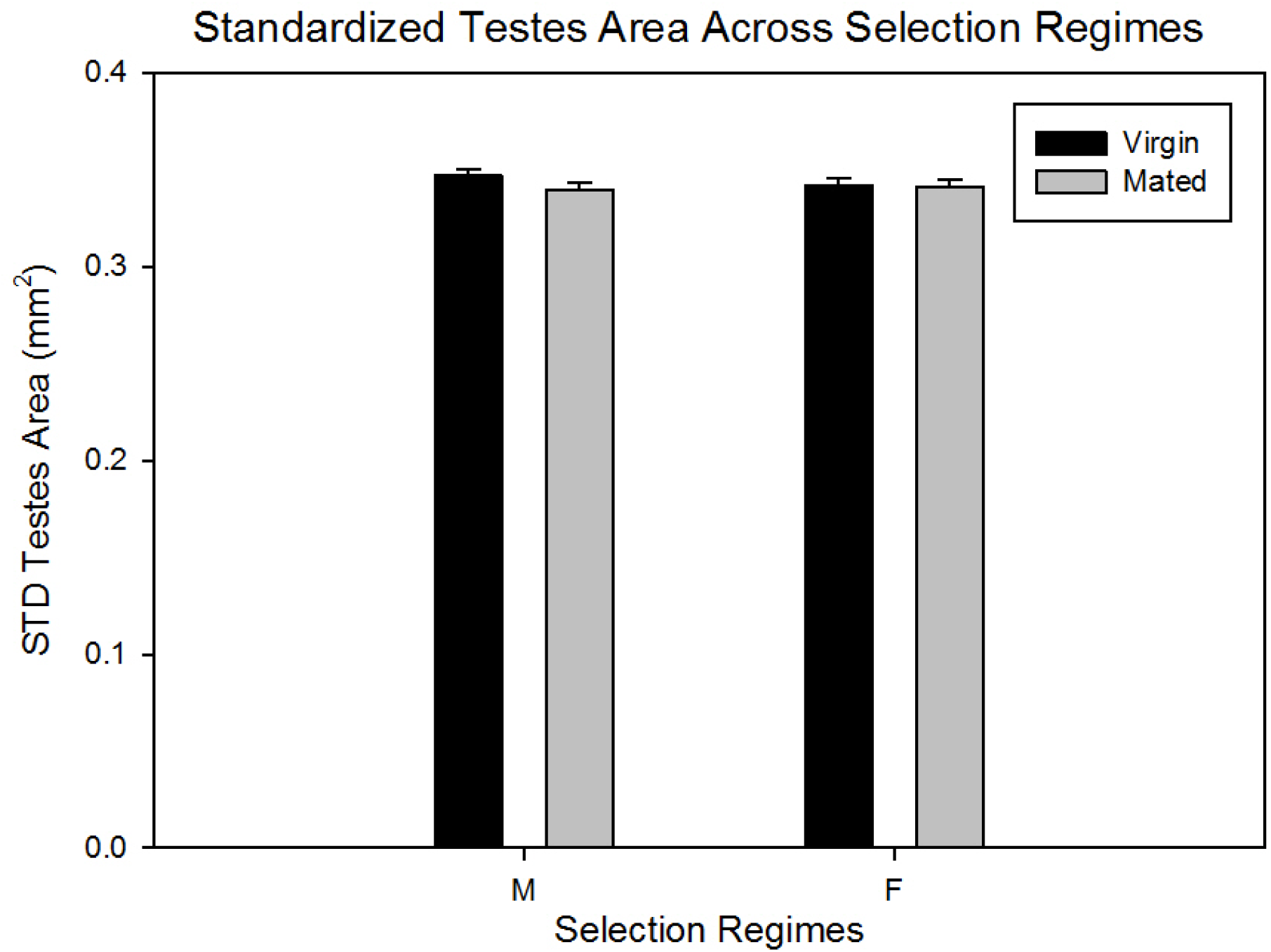
Effect of selection regime and mating status on testes cross section area.

We did not find any significant effect of selection on accessory gland area as well. (Table 2). Mating status had a significant effect on accessory gland size. Mated males from both M and F regime had significantly smaller accessory gland area as compared to virgin males (Figure 7). However, we did not find any significant Selection regime × mating status interaction (Table 1.) indicating that the M and F males are not significantly different in terms of the amount of ejaculate transferred during a single mating.

**Figure 7:**
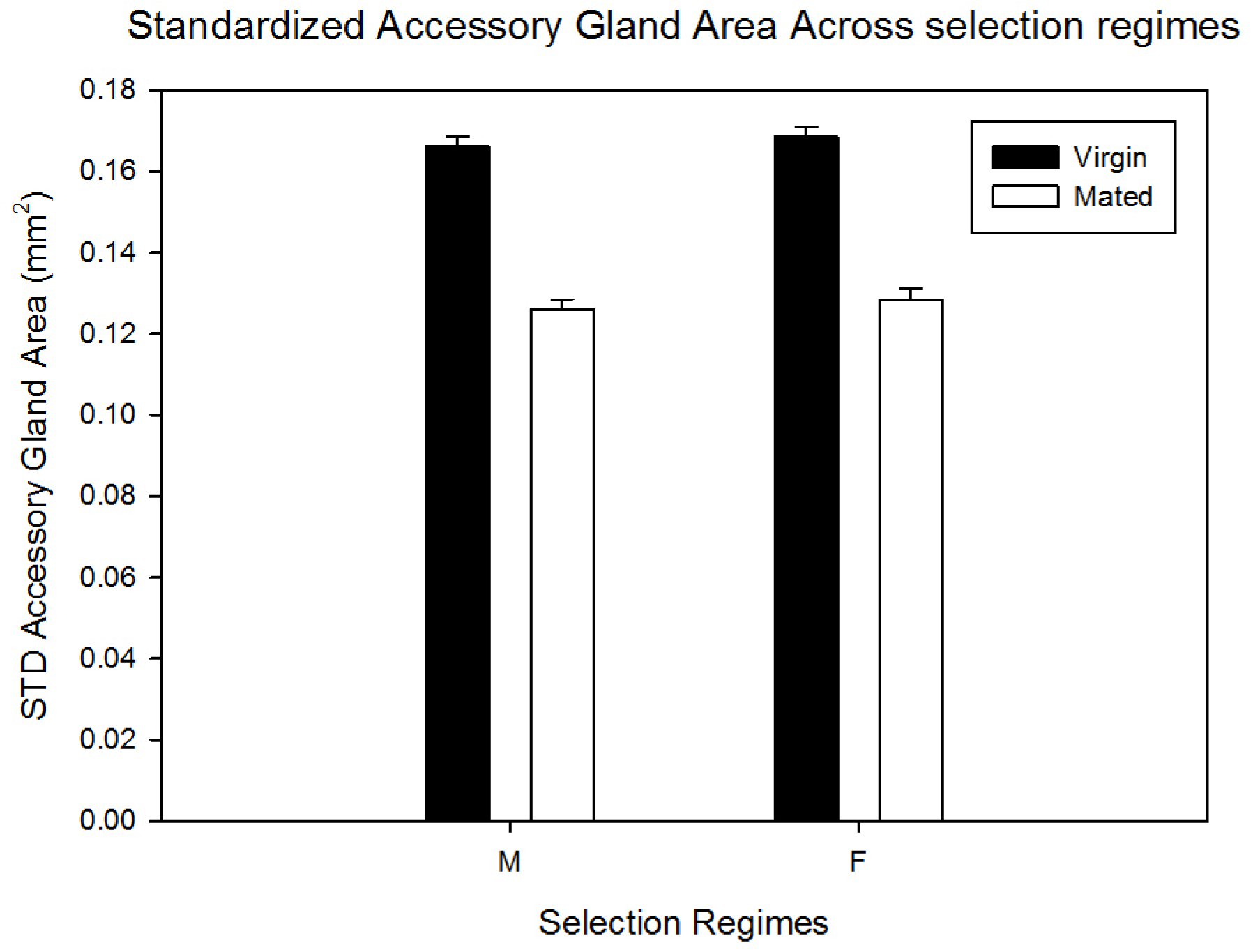
Effect of selection regime and mating status on standardized accessory gland cross section area.

**Table 2:** Multivariabe mixed model analysis of variance (ANOVA) on accessory gland cross section area, with selection regimes and mating treatments as fixed factors and block as random factor.

| Soruce | df num | df den | f-ratio | p-value |
| --- | --- | --- | --- | --- |
| Sel | 1 | 2 | 0.3 | 0.639 |
| Mating Status | 1 | 2 | 25.05 | 0.038 |
| Block | 2 | 2.5 | 5.51 | 0.123 |
| Sel*Mating Status | 1 | 2 | 0 | 0.99 |
| Sel*Block | 2 | 2 | 2.25 | 0.308 |
| Mating Status*Block | 2 | 2 | 7.29 | 0.121 |
| Sel*Mating Status*Block | 2 | 287 | 1.42 | 0.244 |

## Discussion

(Nandy et al., 2013) showed that altering operational sex ratios leads to the evolution of sperm competitive ability - M males evolve increased sperm defense as well as sperm offense ability compared to the F males. Therefore it is quite evident that intensity of sperm competition is different in these two regimes. Our study shows that in investment in reproductive tissue, i.e., testis and accessory gland does not necessarily evolve as a result of change in the risk and intensity of post-copulatory sexual selection-imposed here by altering operational sex-ratio. Neither does investment in ejaculate change over a single mating.

Testis area was not significantly different between virgin M and F males. Thus, contrary to theoretical expectation (Parker, 1970a) and observations from other studies (Hosken & Ward, 2001), altered levels of sperm competition have not lead to the evolution of testis size in our populations even after more than 140 generations of selection. The difference of our result from that of (Hosken & Ward, 2001) could be explained by the difference in the intensity of selection; however it is unlikely as they found difference in testis size after only 10 generations of selection. An alternative explanation is that may be absolute sperm number is more important in *Scatophaga stercoraria* than in *D. melanogaster* (Parker, 1970b).Also, These results are consistent with those of previous studies in *D. melanogaster* (Bhangam et al., 2002) (Linklater et al., 2007).

In case of *Drosophila melanogaster*, accessory gland proteins play a major role in determining the outcome of sperm competition (Frank W. Avila et al., 2012). Therefore, besides testis size (as in Hosken & Ward, 2001) accessory gland size may also evolve in response to increased sperm competition. We did not find any significant difference between the standardized accessory gland area of virgin M and F males, suggesting that investment in accessory glands has also not evolved under different levels of sperm competition. This also is in accordance with the similar previous study by (Linklater et al., 2007).

As mentioned before, another way of evolution under increased sperm competition is increased ejaculate investment in mating. This was measured by reduction in testis and/or accessory gland. We did not find any significant reduction in testis size upon mating. Accessory gland size was affected by mating, with the accessory gland size of mated males being significantly smaller than that of the virgin males. But, the change is accessory gland size after a single mating was not significantly different across selection regimes. Thus, it is unlikely that the amount of Acps transferred by M and F males in a single mating is significantly different. This result is in contrast to that of a previous study in which, males from the male biased (MB) populations showed a faster Acp depletion pattern compared to males from female biased (FB) populations (Linklater et al., 2007). They found that testis size decrease as well upon mating, albeit not differently between MB and FB populations.

(Linklater et al., 2007) assayed change in testes/accessory gland size after five consecutive mating by the males, each with a virgin female. This is likely to represent a very unrealistic scenario, for (Linklater et al., 2007) themselves suggest that even after significant overestimation, the MB males mate 2.68 times on an average. Thus in the MB regime it is safe to assume that males never mate 5 times with virgin females. Further, to the best of our knowledge, there is no clear study showing evolution of sperm competitive ability in those populations. Therefore the biological significance of a difference in accessory gland size after five consecutive mating in the context of the selection regime, is not clear. We measured depletion after a single mating with a virgin female which probably represents the highest amount of ejaculate investment by a male in a given copulation (Ball, M. A. & Parker, G. A., 2007). With the valid assumption that M males have fewer mating opportunities than F males, it was therefore possible that an M male would transfer more ejaculate as and when it has an opportunity to mate. Thus our study’s primary goal was to assess whether increased sperm competitive ability in M males were connected to increased depletion of sperm and/or accessory gland proteins.

One possibility as to why there is no difference in size or depletion of two of the major reproductive organs is that rather than their total quantity, the quality of sperms and/or Acps might have evolved in our populations. There are different indicators of sperm quality such as motility and size. (Gage, 1994) found that across 74 butterflies’ species, sperm length increases with increased sperm competition. Similarly, it is quite possible that sperm size or motility might have evolved in our populations. Alternatively (or additionally), the M and F populations might have evolved with respect to the quality of their Acps. There is indirect evidence supporting this proposition. Using the same populations, (Nandy et al., 2013) found that the females mated (just once) to F males laid significantly more number of eggs compared to females mated to M males. It is very likely that the effect of M and F males on female fecundity is mediated through differences in the quality of the ejaculate, particularly, the Acps, e.g., ovulin or sex peptide. It’s also possible that relative titre of specific ACPs have changed instead of total quantity, a hypothesis that needs further attention.

In conclusion, our study indicates that an evolutionary change in sperm competitive environment does not always necessitate either a change in investment in reproductive tissue or a change in ejaculate quantity or depletion. We propose that when it comes to sperm competition quality can be as important (if not more) quantity, which needs to be given more attention.

## Acknowledgement

We thank Indian Institute of Science Education and Research Mohali, for funding this study.References

